# A Framework of All Discovered Immunological Pathways and Their Roles for Four Specific Types of Pathogens and Hypersensitivities

**DOI:** 10.1101/006965

**Authors:** Wan-Chung Hu

**Affiliations:** Department of Clinical Pathology, Far Eastern Memorial Hospital, New Taipei City, Taiwan; Department of Clinical Pathology, Taipei Tzu Chi Hospital, New Taipei City, Taiwan

**Author notes:** **Correspondence:** Wan-Chung Hu, (W. Hu).

**Keywords:** Th1/2, Th17, Th9, Th22, Treg, Tfh

## Abstract

Tfh initiates four eradicable immunities. Tfh includes FDC, LTi, IL21 CD4 T cell, and IgG/M B cell. Treg initiates four tolerable immunities. Treg includes DCreg, ILCreg, TGFβ CD4 T cell, and IgA B cell. TH1/TH1-like is immunity for intracellular bacteria/protozoa and type 4 delayed type hypersensitivity. TH1 includes M1 macrophage, mDC2, Tc1 CD8 T cell, IFNg CD4 T cell, ILC1, iNKT1, and IgG3 B cell. TH1-like includes M2 macrophage, ILC1, suppressive CD8 T cell, IFNg/TGFβ CD4 T cell, regulatory iNKT cells, and IgA1 B cell. TH2/TH9 is immunity for helminths and type1 IgE mediated hypersensitivity. TH2 includes iEOS eosinophil, Langerhans cell, basophil/MCt mast cell, IL-4 CD4 T cell, ILC2, iNKT2, and IgE/IgG4 B cell. TH9 includes rEOS eosinophil, basophils/mast cell MCct, IL-9 CD4 T cell, ILC2, regulatory iNKT cells, and IgA2 B cell. TH22/TH17 is immunity for extracellular bacteria/fungi and type 3 immune complex hypersensitivity. TH22 includes N1 neutrophils, mDC1, IL-22 CD4 T cell, ILC3(NCR+), iNKT17, and IgG2 B cell. TH17 includes N2 neutrophils, IL-17 CD4 T cell, regulatory iNKT cells, ILC3(NCR−), and IgA2 B cell. THαβ/TH3 is immunity for viruses and type 2 antibody dependent cytotoxic hypersensitivity. THαβ includes NK1 natural killer cell, pDC, Tc2 CD8 T cell, IL10 CD4 T cell, ILC10, iNKT10, and IgG1 B cell. TH3 includes NK2 natural killer cell, suppressive CD8 T cell, ILC10, IL-10/TGFβ CD4 T cell, regulatory iNKT cells, and IgA1 B cell.

**Summary sentence:** The summarized framework of host immunities to explain their relations to specific pathogens and hypersensitivities

## Background of host immunities

Many host immunological pathways have been discovered, such as traditional TH1/TH2, TH3, TH17, TH22, Tfh, Treg, TH9, and Tr1 (THαβ). These identified pathways are not logically organized. In this study, I propose a detailed and complete picture regarding host immunological pathways (Fig. 1).

**Fig 1.**
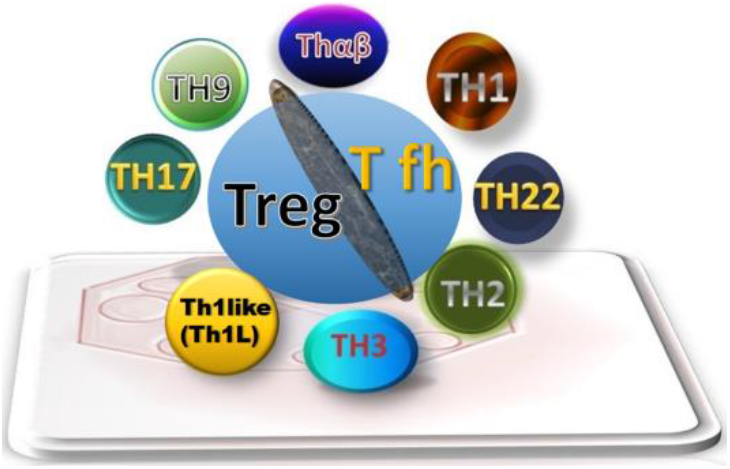
Summary diagram of host immunological pathways. In the middle, the Tfh side (follicular help T cell) initiates initiatory immunity and the Treg side (regulatory T cells) initiates regulatory immunity. Eradicable TH1 and tolerable TH1-like (Th1L), eradicable TH2 and tolerable TH9, eradicable TH22 and tolerable TH17, and eradicable THαβ and tolerable TH3 are related along diagonal lines.

The traditional TH1/TH2 paradigm was proposed by Dr. Mosmann in 1986 ^1^. TH1 was thought to provide host immunity against viruses and intracellular bacteria. TH2 provides host immunity against multicellular parasites (helminthes). In my PhD thesis, I proposed a new THαβ immunological pathway against viruses derived from traditional TH1 immunity^2^. TH1 immunity focuses on intracellular bacteria and protozoa. TH3 immunity and Tr1 immunological pathways were identified after TH1 and TH2 ^3,4^. Recently, additional immune responses have been discovered, including TH17, TH22, Tfh, Treg, and TH1-like immunological pathways ^5–7^.

### Initiatory immune response

Follicular helper T cells (Tfh) are believed to be key helper cells for B-cell germinal centers in lymph nodes. Tfh cells are characterized by IL-21-producing T cells ^8^. Follicular dendritic cells (CD14+) are the antigen presenting cells^9^. Lymphoid tissue inducer cells (LTi) are the innate lymphoid cells for Tfh^10^. BCL6 is a key transcription factor for Tfh. TGFβ with an STAT5 signal can constrain the differentiation of IL-21-producing helper T cells ^11^. IL-21 production is related to STAT1and STAT3 activation. IL-21 production is also related to STAT5 activation because immunosuppressive prolactin can cause STAT5a to suppress BCL6 expression ^12^. However, STAT5b can upregulate BCL6^13^. STAT5a and STAT5b have distinct target genes in immune responses ^14^. The transcription factor that induces Tfh should be STAT5b. BCL6 is important in Tfh development ^15^. Tfh can induce B-cells to start producing the IgM antibody and IL-21 also helps to B cell isotype switch to IgG^16,17^. Besides protein antigen recognized by B cells and T cells, lipid antigens also are recognized by NKT cells. The subtype iNKTfh plays a role in immunity of follicular helper T cells^18^. Thus, T lymphocytes are the first to initiate adaptive host immunity ^19–21^. Different STAT proteins regulate different immunological pathways. If the infection tends to be eradicable, the host immunological pathways mentioned in the following text are generated with other cytokines.

### Eradicable immune responses

TH1 immunity is driven by IL-12 and is the host immunity against intracellular bacteria or protozoa. The antigen presenting cells for TH1 immunity are type 2 myeloid dendritic cells (CD141+ mDC2)^9^. The main effector cells of TH1 immunity are stimulatory macrophages (M1), IFNg-secreting cytotoxic CD8 T cells (CD28+ Tc1), IFNg-secreting CD4 T cells, iNKT1 cells, and IgG3-producing B-cells ^22,23^. The initiation of eradicable immunity also needs the help of innate lymphoid cells to produce initial cytokines to drive to different immunological pathway. For TH1 immunity, the key innate lymphoid cells are ILC1^24^. The key transcription factor for TH1 immunity is STAT4. T-bet also plays a vital role in the TH1 immunological pathway. TH1 immunity against self-antigen is type 4 delayed-type hypersensitivity, such as type 1 diabetes mellitus.

TH2 immunity is driven by IL-4. TH2 provides immunity against extracellular parasites (helminths). The antigen presenting cells for TH2 immunity are Langerhans cells (CD1a+)^9,25^. The main effector cells of TH2 immunity are eosinophils (iEOS), basophils/proinflammatory tissue mast cells (MCt, mast cell tryptase), IL-4/IL-5-secreting CD4 T cells, iNKT2 cells, ILC2, and IgG4/IgE-producing B-cells ^26^. IgG4 activates eosinophils, and IgE activates mast cells, such as in acute anaphylaxis ^27^. The function of IgG4–eosinophil is to activate eosinophil-mediated cellular immunity against parasites or insects. The function of IgE–mast cells is to expel helminths or insects through a physiological mechanism. Mast cells activated by IgE can release histamine, which causes bronchoconstriction, vomiting/nausea, rhinorrhea, skin itchiness, stomach acidification, increased local vascular permeability, or increased bowel movement. These actions can all help to expel helminths or insects physiologically. The key transcription factor for TH2 immunity is STAT6. GATA3 also plays a vital role in the TH2 immunological pathway. TH2 immunity against self-antigen is a type1 immediate allergy, such as food/drug allergy or urticaria.

THαβ is distinguished from traditional TH1 immunity ^2^. THαβ immunity is against viruses and was called Tr1 cell by a few previous researchers^4^. THαβ immunity is driven by IFNa/b or IL-10. The antigen presenting cells for THαβ immunity are plasmacytoid dendritic cells (pDC)^9^. The main effector cells of THαβ immunity are IL-10-producing stimulatory NK cells (CD56−CD16 + NK1 cells), IL-10/IL-27-secreting CD4 T cells, IL-10-secreting cytotoxic CD8 T cells (CD28+ Tc2), iNKT10 cells, ILC10, and IgG1-producing B-cells ^22,28–30^. The CD27 molecule is important for virus immunity. The key transcription factors for THαβ immunity are STAT1and STAT2^31^. THαβ immunity against self-antigen is a type 2 antibody-dependent cytotoxic hypersensitivity, such as an acute stage of myasthenia gravis. IL-10 is not merely an immunosuppressive cytokine; it can also have potent stimulatory effects on NK cells, CTLs, and B-cells.

TH22 is the host innate immunity against extracellular bacteria and fungi. TH22 is driven by IL-6 or TNFa ^32^. The antigen presenting cells for TH22 immunity are type 1 myeloid dendritic cells (CD1c+ mDC1)^9^. The main effector cells for TH22 immunity are PMNs (N1), IL-22-secreting CD4 T cells, complements, pentraxins, iNKT17 cells, ILC3(NCR+), and IgG2-producing B-cells ^6,33^. The key transcription factor for TH22 is STAT3. AP1 and CEBP are also important transcription factors. TGFβ can suppress IL-22 to skew to TH17 immunity ^34^. TH22 against self-antigen is a type 3 immune-complex and complement-mediated hypersensitivity, such as the Arthus reaction. The extracellular or intracellular locations of protozoa or fungi mainly decide the host immunological pathways.

Four IgG subtypes fit the four types of acute immunological pathways. Murine IgG antibodies also have four subclasses. The following correlation exists between murine and human IgG subtypes: Human IgG1<->Murine IgG2a; Human IgG2<->Murine IgG3; Human IgG3<->Murine IgG2b; and Human IgG4<->Murine IgG1 ^35^. hIgG1/mIgG2a works against viral antigens; hIgG2/mIgG3 works against bacterial antigen, especially polysaccharides; hIgG3/mIgG2b works against intracellular bacteria; and hIgG4/mIgG1 works against parasite antigens ^36–38^. It is also worth noting that the immune response against fungus or protozoa is mainly based on intracellular or extracellular. Extracellular fungus such as Candida spp or Aspergillus spp usually trigger TH22 immunity, but intracellular fungus such as Histoplasma spp will trigger TH1 immunity.

### Regulatory immune responses

Treg is the host immune inhibitory mechanism ^39^ and is driven by IL-2 and TGFβ. Regulatory dendritic cells (DCreg) are the antigen presenting cells for Treg^40^. Regulatory innate lymphoid cells (ILCreg)are the initial helpers for Treg production^41^. The main effector cells for Treg are the TGFβ-producing CD4 T cells, FOXP3 regulatory iNKT cells, and IgA-producing B-cells^42^. The key transcription factor for the Treg pathway is STAT5, especially STAT5a. However, both STAT5a and STAT5b play nonredundant roles in Treg generation ^43^. They may act sequentially with STAT5b activation first in Tfh signaling. STAT5b and STAT5a combined signaling induces Treg generation. The combination of Treg and the aforementioned four immunological pathways is important to shift adaptive immunity to tolerable immunity. During initial infection, acute-stage fierce cytokines can rapidly kill pathogens and infected cells or tissues. However, if the pathogen infects numerous cells in a tissue, such as the liver, killing the infected cells would completely destroy the organ ^44^. Thus, a regulatory T cell STAT5 signal combined with TH1/TH2/TH22/THαβ would allow the generation of CD4 T cells with less fierce cytokines ^43^. TH1-like/TH9/TH17/TH3 immunological pathways are generated during chronic infection. IgA1 and IgA2 are the two types of IgA antibodies. IgA1 is the dominant IgA antibody in serum, whereas IgA2 is the dominant IgA in mucosa. TGFβ can induce IgA1 or IgA2 depending on the lymphoid follicle location ^45^. In GULTs or Peyer’s patch, IgA2 is the dominant IgA antibody produced in the GI mucosa. In the lymph nodes of other body locations, IgA1 is the dominant IgA antibody produced. However, IgA1 is especially related to viral protein antigens, whereas IgA2 is especially related to bacterial antigens, such as LPS. The heavy-chain locus sequence of B-cell antibodies is IgM, IgD, IgG3, IgG1, IgA1, IgG2, IgG4, IgE, and IgA2. B-cells double-express IgM and IgD. IgG3, IgG1, and IgA1 comprise the first group for cellular immunity. IgG2, IgG4, IgE, and IgA2 can be viewed as the second group for humoral immunity. The gene sequence order is important, and it affects the time sequence of the isotype switch. IL-13 is a profibrogenic Treg-related cytokine related to TGFβ signaling.

### Tolerable immune responses

TH1-like cells (non-classic TH1) are initiated by TGFβ (STAT5 signaling) and IFNg (STAT4 signaling). TH1-like cells with Foxp3 + regulatory character have been identified ^7^. TH1 helper cells and TH1-like cells are closely related ^46^. TH1-like cells are chronic host immunity of the TH1 immune response. Thus, these cells could be related to chronic inflammation, such as long-term tuberculosis infection. The effector cells of TH1-like immunity include suppressive macrophages (M2), ILC1, suppressive CD8 T cells (CD28-CD8+Treg), IgA1-producing B-cells, regulatory iNKT cells, and IFNg/TGFβ-producing CD4 T cells ^23,33,47^. TH1-like immunity induces type 4 delayed-type hypersensitivity, such as Crohn’s disease.

TH9 cells are driven by IL-4 (STAT6 signaling) combined with TGFβ (STAT5 signaling) ^48–50^. Thus, TH9 cells are closely related to the TH2 immunological pathway. The cells are characterized by the IL-9-secreting CD4 T cell. TH9 cells are important under a chronic allergic condition, such as asthma. Thus, TH9 helper cells are chronic T helper cells related to TH2 immunity. The effector cells of TH9 immunity include regulatory eosinophils, basophils/profibrotic mast cells (MCct, mast cell chymase & tryptase), ILC2, IL-9-producing CD4 T cells, regulatory iNKT cells, and IgA2-producing B-cells ^33,51^. TH9 immunity induces type1 allergy, including asthma ^26^.

TH17 cells are driven by IL-6/IL-1 combined with TGFβ ^5^. Thus, TH17 cells are closely related to the TH22 immunological pathway. TH17 cells are characterized by the IL-17-secreting CD4 T cell. TH17 cells are important in chronic immune-complex-mediated diseases, such as rheumatic arthritis. The TH17 helper cell is the chronic T helper cell related to TH22 immunity. TGFβ with STAT5 can suppress the acute IL-22-producing cells and enhance the chronic IL-17-producing cells ^34^. Because of the role of TGFβ in TH17 immunity, regulatory IL-17-producing cells are noted. The effector cells of TH17 immunity include regulatory neutrophils (N2), ILC3(NCR−), IL-17-producing CD4 T cells, regulatory iNKT cells, and IgA2-producing B-cells ^33,52^. TH17 immunity induces type 3 immune-complex hypersensitivity, including ulcerative colitis.

TH3 cells are driven by IL-10 and TGFβ ^53,54^. Thus, TH3 cells are closely related to the THαβ immunological pathway. These cells also produce IL-10 and TGFβ. Thus, TH3 helper cells are important to chronic antibody-dependent cellular cytotoxic hypersensitivity. TH3 cells are the chronic helper T cells corresponding to the THαβ helper cells. The TH3 immune effector cells include IL-13-producing regulatory NK cells (CD56 + CD16−NK2 cells), ILC10, IL-10- and TGFβ-secreting CD4 T cells, suppressive CD8 T cells (CD28-CD8+ Treg), regulatory iNKT cells, and IgA1-producing B-cells ^29,30,47,55,56^. IgA1 is produced in serum and works against viral protein antigens. TH3 immunity induces type 2 antibody-dependent cytotoxic hypersensitivity, including a chronic stage of SLE.

### Concluding remarks

The summary diagram includes complete pictures of the 4 × 2 + 2 immunological pathways (Table 1). The TH1, TH2, TH22, and THαβ eradicable immune responses correspond to the TH1-like, TH9, TH17, and TH3 tolerable immune responses, respectively, and match the four types of hypersensitivity. The detailed immune response against different pathogens and allergy/hypersensitivity can be clearly understood.

**Table 1.**
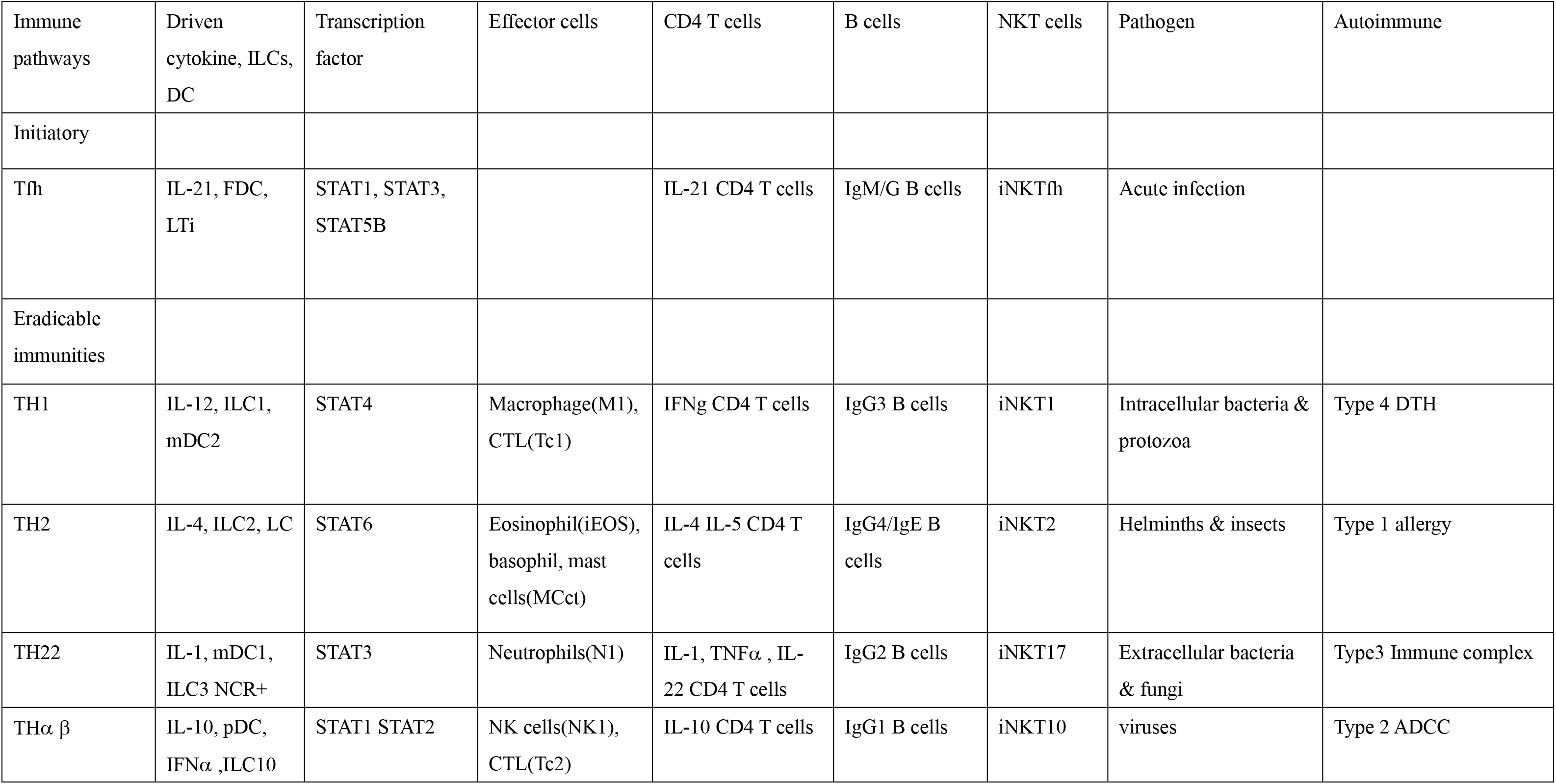
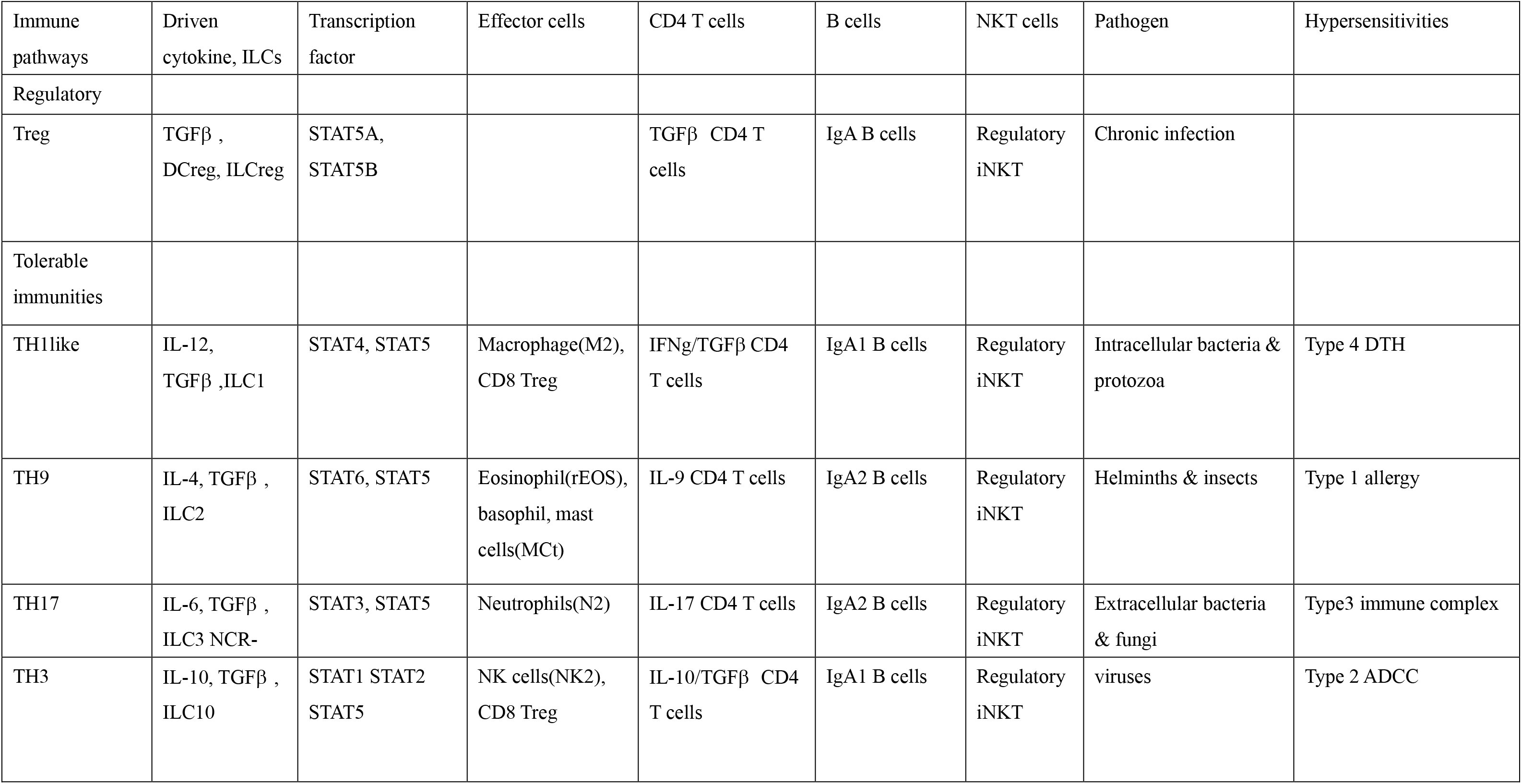
Summary of host immunological pathways

## Authorship

The author, Wan-Chung Hu, fully contributes to the concept, literate search, writing, and final approval of this manuscript.

## Acknowledgment

The author is very thankful for the instructions provided by Professors Alan Scott, Louis August Bourgeois, and Pien-Chien Huang during his PhD study on vaccine science from the Department of International Health, Johns Hopkins University, Bloomberg School of Public Health. The author is also very grateful for the guidance of Professors Chi-Huey Wong and Alice Lin-Tsing Yu in his postdoctorate research in the Genomics Research Center of Academia Sinica, Taiwan.

## Author’s information

The author, Wan-Chung Hu, graduated as a MD from National Taiwan University. He then completed his PhD in vaccine science from the Department of International Health, Johns Hopkins University, Bloomberg School of Public Health. The author’s PhD thesis was on host immune reaction against malarial infection. He conducted a postdoctorate study on cancer immunotherapy at the Genomics Research Center, Academia Sinica, Taiwan. The author completed his PGY training from Mackay Memorial Hospital and Shin-Kong Memorial Hospital in Taiwan. He is currently working as a chief resident in the Department of Clinical Pathology, Far Eastern Memorial Hospital, Taiwan (R.O.C.), for resident training.

Please see the brief abstract in **BioRxiv doi:** https://doi.org/10.1101/006965.

